# Proximity Labeling-Assisted Flow Cytometry Improves Fluorescence Contrast in Extracellular Vesicle-Enriched Fractions

**DOI:** 10.64898/2026.09.15.751677

**Authors:** Leona Kashimata, Kensuke Iwasa, Megumi Kumagai, Ryoko Sasaki, Shun Shinomiya, Machika Soma, Hidetoshi Iemura, Ryu Sekiya, Isaku Horiuchi, Masahito Katsuki, Kumiko Komatsu, Yosuke Mizuno, Hidetoshi Nakamura, Makoto Nagata, Tsuyoshi Sato, Ko Ito, Yohei Kawasaki, Kazuyuki Nakagome, Norihiro Kotani

## Abstract

Extracellular vesicles (EVs) are potential sources of circulating biomarkers, but disease-associated EVs are often scarce and their nanoscale dimensions limit analysis by conventional flow cytometry. To improve the detection of disease-associated EV signals using widely available instruments, we developed a workflow combining preparation of serum EV-enriched fractions by size-exclusion chromatography, CHL1-initiated enzyme-mediated activation of radical sources (EMARS) proximity labeling, and conventional flow cytometry. EMARS deposits fluorescein on molecules proximal to antibody-bound CHL1, thereby increasing target-initiated fluorescence. In a representative comparison, EMARS increased the signal-to-background ratio from 1.21 with conventional antibody labeling to 6.28. Because unlabeled EVs and debris interfered with analysis, we used an AI-assisted procedure to explore FITC-H cutoffs for retained P1 events. ChatGPT compared 65 candidate cutoffs across statistical and event-retention metrics and recommended 1000; the authors reviewed the complete results before adopting this value. At this data-selected cutoff, mean fluorescence intensity was higher in interstitial lung disease (ILD) than in non-ILD samples among 28 evaluable samples (P = 0.002; area under the receiver operating characteristic curve [AUC] = 0.846); neither value was adjusted for cutoff selection. Post hoc nested leave-one-out cross-validation (LOOCV), in which cutoff selection was repeated within each outer fold, yielded an AUC of 0.716 with 29 of 34 samples evaluable. The association with ILD requires validation in independent cohorts, but these exploratory findings suggest that proximity labeling may enhance measurable fluorescence contrast in serum EV-enriched fractions and support further evaluation of disease-associated EV signals using conventional flow cytometers.

## 1. Introduction

Extracellular vesicles (EVs) are lipid bilayer-delimited particles released by cells and are present in a variety of body fluids [1,2]. EVs contain proteins, lipids, and nucleic acids derived from their cells of origin and have therefore been investigated as mediators of intercellular communication and as potential biomarkers [3]. Because EVs are present in readily accessible body fluids and carry molecular information related to their cells of origin, analytical methods that can be implemented in general clinical and research laboratories are important for translating EV research into biomarker studies [3,4].

Among the available methods for EV analysis, flow cytometry enables rapid, multiparametric analysis of large numbers of samples [4]. High-sensitivity flow cytometry platforms have enabled surface-protein analysis at the single-EV level [5]. However, access to these platforms varies among institutions. Improving the detection of disease-associated EV signals using more widely available conventional flow cytometers may broaden the settings in which EV-based clinical testing can be performed [4].

Conventional flow cytometers are primarily optimized for cell analysis rather than nanoscale particles [4]. Because EVs are substantially smaller than cells, their scatter and fluorescence signals may be near or below the detection limits of these instruments [4,6–8]. Coincidence or swarm detection may cause multiple particles to be detected as a single event [4,6,9], and fluorescence from antibodies bound to target molecules may be difficult to distinguish from background signals arising from the instrument and sample buffer. Antibody aggregates and unbound antibodies can also contribute to false-positive fluorescence events [4,6,10]. The present study addressed the limited intensity of target-associated fluorescence relative to the background against which it is recorded, rather than the scatter sensitivity of these instruments. To mitigate these limitations as far as possible, we focused on an approach to increase the fluorescence intensity of target EVs relative to the background.

Enzyme-mediated activation of radical sources (EMARS) is a proximity labeling method that uses horseradish peroxidase (HRP) conjugated to a target-binding molecule [11]. EMARS has been used to identify molecular complexes on cell membranes [12,13] and to analyze EV surface proteins [14,15]. In the presence of hydrogen peroxide, HRP activates fluorescein-tyramide to generate short-lived radicals. Because of their short lifetime, these radicals react with molecules in the immediate proximity of the enzyme and covalently attach fluorescein to them [11,15]. By depositing fluorescein around an antibody-bound target, EMARS may amplify target-associated fluorescence beyond that obtained by direct antibody labeling.

We hypothesized that EMARS-mediated signal amplification would permit EV surface antigen-targeted EV-associated fluorescence to be measured using conventional flow cytometry. We therefore sought to develop an analytical workflow combining size-exclusion chromatography (SEC)-based preparation of serum EV-enriched fractions, EMARS labeling, and measurement using a typical flow cytometer.

In this study, close homolog of L1 (CHL1), which has previously been used as a surface target for EVs, was selected as a model target [14,15]. We attempted to label and specifically detect CHL1-expressing EVs using EMARS; however, detection remained challenging because of issues such as background signals. We therefore sought to address these limitations by establishing operational flow-cytometry gating parameters. Gating for side-scatter (SSC) signals was set using commercially available nanoparticle standards. In contrast, for fluorescence-intensity gating, we compared candidate FITC-H thresholds using clinical serum samples from patients with pulmonary disease. Comparing candidate gating parameters required testing numerous configurations and calculating statistical measures and the area under the receiver operating characteristic curve (AUC) for each configuration. Accordingly, we developed an AI-assisted scheme to automate these calculations and comparisons. As an exploratory application, the workflow was applied to clinical serum samples, in which MFI was unexpectedly higher in samples from patients with interstitial lung disease (ILD) than in samples from patients without ILD.

## 2. Materials and methods

### 2.1. Patient serum samples

Human serum samples were obtained from patients evaluated for respiratory diseases in the Department of Respiratory Medicine, Saitama Medical University Hospital (Table S1A and S1B). The study was conducted in accordance with the Declaration of Helsinki and was approved by the Institutional Review Board of Saitama Medical University Hospital (approval no. HP2021-067). Written informed consent for participation in the study was obtained from all participants.

Clinical samples were classified into three groups: patients with interstitial lung disease (ILD) without an active concurrent thoracic malignancy, patients with a thoracic malignancy without ILD, and patients with other respiratory diseases without ILD. The latter two groups were pooled as the non-ILD group. The primary analysis included 34 samples: 18 in the ILD group and 16 in the non-ILD group, comprising 11 samples from patients with thoracic malignancies and five from patients with other respiratory diseases.

Three samples that could not be assigned unambiguously to either comparison group were excluded from the primary analysis. Sample 5 was obtained from a patient with ILD and recurrent lung adenocarcinoma, and Sample 92 was obtained from a patient with ILD and malignant pleural mesothelioma; both samples therefore met the definitions of both groups. For Sample 102, neither the suspected interstitial lung involvement associated with IgG4-related disease nor the nature of the mediastinal lymphadenopathy had been established at the time of classification. These three samples were included in the ILD group in a sensitivity analysis. Clinical classification was based on finalized clinical information. Patient characteristics are summarized in Table 1.

**Table 1.** Clinical characteristics of the primary analysis set (n = 34).

| Characteristic | ILD | Non-ILD, cancer | Non-ILD, other |
| --- | --- | --- | --- |
| No. of patients | 18 | 11 | 5 |
| Age, years, median (range) | 74 (61-82) | 70 (52-82) | 75 (48-81) |
| Male sex, n (%) | 13 (72) | 7 (64) | 4 (80) |
| Smoking status, n (%) |  |  |  |
| Never | 3 (17) | 4 (36) | 2 (40) |
| Ever | 15 (83) | 7 (64) | 3 (60) |
| Brinkman index, median (range)<sup>a</sup> | 500 (0-1320) | 800 (0-2320) | 300 (0-1320) |
| Thoracic malignancy at sampling, n (%) | 0 (0) | 11 (100) | 0 (0) |
| Extrathoracic malignancy, n (%)<sup>b</sup> | 4 (22) | 0 (0) | 1 (20) |
| KL-6, U/mL, median (range)<sup>c</sup> | 868 (197-2571) | 434 (222-1344) | 148 |

ILD, interstitial lung disease; KL-6, Krebs von den Lungen-6. Each sample was obtained from a different patient. Continuous variables are presented as median (range) and categorical variables as n (%). Characteristics are descriptive; no between-group comparisons were performed. Three samples (Sample IDs 5, 92, and 102) were excluded from the primary analysis.
<sup>a</sup> Brinkman index (cigarettes per day multiplied by years of smoking) was calculable in 17, 11, and 5 patients in the ILD, cancer, and other groups, respectively; never-smokers were assigned a value of 0.
<sup>b</sup> Extrathoracic malignancy included current or previous malignancies identified during clinical review; one hematologic malignancy was diagnosed 63 days after sample collection.
<sup>c</sup> KL-6 values obtained within 30 days of blood sampling were included; measurements were available in 10, 5, and 1 patients, respectively, and were obtained as part of routine clinical care rather than systematically for this study.

Serum aliquots were stored at −80 °C and subjected to no more than one freeze-thaw cycle. After thawing, the samples were kept at 4 °C and generally processed on the same day. Before SEC, the serum was centrifuged at approximately 840 × g (3000 rpm) for 5 min at 4 °C using a Centrifuge 5417R equipped with an FA-45-24-11 fixed-angle rotor (Eppendorf, Hamburg, Germany). The supernatant was used in the subsequent procedures.

### 2.2. Preparation of serum EV-enriched fractions by SEC

Serum EV-enriched fractions were prepared using a modification of the method described by Kaneda et al. [14]. Sephacryl S-500 High Resolution resin (catalog no. 17061310; Cytiva Sweden AB, Uppsala, Sweden) was pretreated for 1 h with a yeast extract solution (0.1 g/100 mL H_2_O; BD Biosciences) to reduce nonspecific adsorption of EVs to the resin. A Bio-Spin disposable chromatography column (catalog no. 7326008; Bio-Rad Laboratories, Hercules, CA, USA) was prefilled with 1 mL of phosphate-buffered saline (PBS). After gentle mixing, 1 mL of the resin suspension was gently added to the PBS while avoiding the introduction of air bubbles. The column was then equilibrated with an additional 1 mL of PBS. All subsequent elution was performed by gravity flow.

Clarified serum (50 µL) or EV-free PBS as a negative control (PBS control) was applied to the column. After the sample had entered the resin bed, 250 µL of PBS was added, and the eluate was discarded. A further 150 µL of PBS was then added, and the resulting 150-µL eluate, corresponding to fractions 6-8, was collected as a serum EV-enriched fraction. Each column was assigned to a single clinical sample and was not shared among samples. The same column was used at three distinct stages of the workflow: to obtain the initial serum EV-enriched fraction, to remove unbound antibody after the anti-CHL1 antibody reaction, and to remove residual reagents after the EMARS reaction. Before each reuse, the column was washed with 3 mL of PBS.

### 2.3. Characterization of serum EV-enriched fractions

#### 2.3.1. Nanoparticle tracking analysis

Nanoparticle tracking analysis (NTA) was performed by Egret Lab Co., Ltd. (Tokushima, Japan) on an SEC fraction prepared from a single clinical serum sample. Measurements were performed using a NanoSight NS300 equipped with a 405-nm laser (Malvern Panalytical, Malvern, UK) and NTA software version 3.4 (build 3.4.4). Each sample was diluted 10-fold in PBS. Five recordings of 1498 frames were acquired at 25 frames/s using a camera level of 12, a detection threshold of 5, and a syringe-pump speed setting of 30. The measurement temperature was 24.1 °C. Particle-size distributions were calculated using the manufacturer’s software.

#### 2.3.2. Transmission electron microscopy

SEC-derived serum EV-enriched fractions were applied to hydrophilic Formvar-coated grids (Nisshin EM, Tokyo, Japan), allowed to adsorb for 10 min, and negatively stained with 2% uranyl acetate. After air-drying, the samples were examined using a JEM-1400 transmission electron microscope (JEOL Ltd., Tokyo, Japan) operated at 80 kV.

#### 2.3.3. Western blot analysis

SEC-derived serum EV-enriched fractions that had not been subjected to EMARS labeling were concentrated using Nanosep centrifugal devices equipped with 30K Omega polyethersulfone membranes (catalog no. OD030C34; Pall Corporation, Port Washington, NY, USA). The devices were centrifuged at 8000 × g for 5 min. Material retained on the membrane was recovered in 20 µL of reducing SDS sample buffer prepared from 6× SDS sample buffer (catalog no. 09499-14; Nacalai Tesque, Kyoto, Japan), mixed by gentle vortexing and pipetting, and heated at 100 °C for 5 min. The entire 20-µL sample was subjected to 8% SDS-PAGE at 200 V and 100 mA for 30 min with Rapi running buffer solution (catalog no. 12981-74; Nacalai Tesque). An Immobilon-P polyvinylidene difluoride membrane (Millipore Corporation, Billerica, MA, USA) was activated in methanol for at least 30 s, and proteins were transferred using a semi-dry transfer system at 100 mA for 60 min. Tris-buffered saline containing Tween 20 (TBST) was prepared from Tris Buffered Saline with Tween 20 Tablets, pH 7.6 (catalog no. T9142; Takara Bio Inc., Kusatsu, Shiga, Japan), according to the manufacturer’s instructions. The membrane was blocked in 5% skim milk in TBST for 1 h at room temperature or overnight at 4 °C. Primary antibodies were diluted 1:1000 in 0.5% skim milk in TBST and incubated with the membrane for 1 h at room temperature or overnight at 4 °C. The primary antibodies were anti-CD63 antibody (catalog no. SAB4301607, lot 2452140192; Sigma-Aldrich, St. Louis, MO, USA), rabbit monoclonal anti-integrin alpha 2 antibody (clone EPR5788, catalog no. ab133557; Abcam, Cambridge, UK), and anti-beta-actin antibody (catalog no. 3662-100; BioVision, Milpitas, CA, USA).

After washing with TBST for 5, 10, and 5 min, the membrane was incubated for 1 h at room temperature with horseradish peroxidase (HRP)-conjugated goat anti-rabbit IgG (H+L) (catalog no. W4011; Promega, Madison, WI, USA) diluted 1:5000 in 0.5% skim milk in TBST. The membrane was then washed five times for 5 min each with TBST and incubated for 1 min with Immobilon Western Chemiluminescent HRP Substrate (catalog no. WBKLS0500; Millipore Corporation). Chemiluminescence was detected using a ChemiDoc MP imaging system and Image Lab software (Bio-Rad Laboratories). A549 and 293T cell lysates were used as positive controls for the respective proteins. Uncropped blot images are shown in Fig. S8.

### 2.4. EMARS labeling

The enzyme-mediated activation of radical sources (EMARS) reaction was performed as described by Kaneda et al. and Kotani et al. [11,14–16], with modifications for the analysis of serum EV-enriched fractions by flow cytometry. An anti-human CHL1 antibody (MAB2126; R&D Systems, Minneapolis, MN, USA) was partially reduced and conjugated to HRP using a Peroxidase Labeling Kit-SH (Dojindo Laboratories, Kumamoto, Japan).

A 100-µL aliquot of the 150-µL serum EV-enriched fraction was transferred to a 1.5-mL tube, and 0.5 µL of HRP-conjugated anti-CHL1 antibody was added with gentle mixing. After incubation for 20 min at room temperature, the column assigned to the sample was washed with 3 mL of PBS, and the reaction mixture was reapplied to the column. PBS (250 µL) was added, and the eluate was discarded. The subsequent 150-µL eluate was collected.

The fluorescein-tyramide (FT) reagent was prepared and added to the antibody-treated eluate as described previously [14–16]. The reaction mixture was then gently mixed and incubated for 15 min at room temperature in the dark. The column assigned to the sample was washed with 3 mL of PBS, and 100 µL of the reaction mixture was applied. After the sample had entered the resin bed, 250 µL of PBS was added, and the eluate was discarded. The subsequent 150-µL eluate was collected directly into a flow cytometry tube.

### 2.5. Conventional antibody labeling

For comparison with EMARS labeling, a PBS control and a serum EV-enriched fraction were incubated with an HRP-conjugated anti-human CHL1 antibody at room temperature for 20 min, followed by labeling with an Alexa Fluor 488-conjugated anti-rat IgG secondary antibody (catalog no. 4416; Cell Signaling Technology, Danvers, MA, USA), in place of the FT reagents, at room temperature for 20 min. The labeled samples were measured using the same flow cytometer as that used for the EMARS-labeled samples. Because conventional antibody labeling and EMARS labeling involve different labeling reactions, absolute fluorescence intensities were not compared directly between the two methods. Instead, the serum-to-PBS fluorescence ratio was calculated separately for each method.

### 2.6. Flow cytometry

Samples were analyzed using a FACSCanto II flow cytometer controlled by FACSDiva software version 6.1 (BD Biosciences, San Jose, CA, USA). Detector voltages were set to 700 V for forward scatter (FSC), 550 V for side scatter (SSC), and 620 V for FITC. Parameters were displayed logarithmically during acquisition, and the acquired data were stored as FCS 3.0 files containing linear-scale parameter values.

### 2.7. SSC-H–based operational event gate

A manually defined operational analysis region, designated P1, was established on SSC-H versus FITC-H plots. P1 was positioned with reference to the distributions of the 0.16-, 0.20-, and 0.24-µm Megamix-Plus SSC particles (reference no. 7803; BioCytex, Marseille, France) and to the PBS process control, while excluding the prominent high-SSC-H/low-FITC-H background region. A session-specific P1 was defined from the reference particles and PBS control and applied to serum samples acquired in that session. Ungated FCS 3.0 files were retained for all samples, and events within P1 were used for the primary analysis. To assess the contribution of P1, analyses with and without P1 were compared using the 28 samples that retained events within P1 at FITC-H = 1000 (15 ILD and 13 non-ILD samples). Without P1, all acquired events at or above FITC-H = 1000 were included. MFI, AUC, and LOOCV AUC were calculated under each condition. This matched comparison was conditional on the presence of events within P1, the analysis region under evaluation. To examine this comparison across the full candidate cutoff range, AUCs with and without P1 were also calculated at each cutoff using only samples that retained events under both conditions at that cutoff; therefore, the matched sample set could vary across cutoffs. A supplementary comparison was also performed without restricting the analysis to the samples evaluable under both conditions. This unmatched comparison included 28 evaluable samples when P1 was applied and 34 when P1 was not applied. Because FITC-H = 1000 was selected within the P1-based workflow, the comparison was considered an exploratory assessment conditional on the previously selected cutoff.

### 2.8. AI-assisted selection of the FITC-H cutoff

After manual definition of P1, candidate FITC-H cutoffs were compared with assistance from ChatGPT (OpenAI, San Francisco, CA, USA; accessed July 6, 2026). For clarity, the term “full-dataset analysis” refers to the exploratory cutoff-selection procedure in which all 34 samples in the primary analysis set entered the comparison of candidate cutoffs. At each cutoff, group comparisons and AUCs were calculated using samples with at least one retained event, while the number and proportion of samples with no retained events and the number of retained events were evaluated as operational metrics. ChatGPT generated and executed code to read the P1-gated FCS 3.0 data, apply each cutoff, calculate sample-level MFI, and perform the specified statistical analyses. It then compared the resulting statistical and event-retention metrics and returned a recommended cutoff with its rationale. A total of 65 candidate cutoffs were evaluated: values from 0 to 3000 in increments of 50 and values from 3500 to 5000 in increments of 500. Events with FITC-H values greater than or equal to each cutoff were retained, and the arithmetic MFI was calculated for each sample. If a sample contained no events at or above a given cutoff, a mean value could not be calculated, and that sample was excluded only from the analysis at that cutoff. At each cutoff, ILD and non-ILD samples were compared using the Mann–Whitney U test. Receiver operating characteristic analysis, univariable logistic regression, and fixed-cutoff leave-one-out cross-validation (LOOCV) were performed as described in Section 2.11. Odds ratios were calculated per 100-unit increase in MFI. For fixed-cutoff LOOCV, one sample was omitted, a logistic regression model was fitted using the remaining samples, and the predicted probability for the omitted sample was calculated. This procedure was repeated for every sample. The FITC-H cutoff was not reselected within individual LOOCV iterations. Candidate cutoffs were screened using operational criteria of P < 0.05 in the Mann–Whitney U test, P < 0.01 in the likelihood-ratio test, and AUC ≥ 0.70. For cutoffs meeting these criteria, ChatGPT compared the AUC obtained from the full-dataset analysis, fixed-cutoff LOOCV AUC, difference between these AUCs, numbers of evaluable samples and samples with no retained events, retained-event count, and event-retention rate. ChatGPT recommended FITC-H = 1000 based on the balance among discrimination, fixed-cutoff cross-validated performance, and event retention. After reviewing the analysis conditions, code, cutoff-specific outputs, and recommendation, the authors adopted 1000 as an exploratory, data-selected cutoff. The arithmetic MFI of events within P1 at or above FITC-H = 1000 was used as the sample-level readout. ChatGPT was not used to define P1 or classify individual events. Its role comprised code generation and execution, comparison of the specified outputs across candidate cutoffs, and generation of a recommendation with an accompanying rationale; the authors made the final decision.

As a separate post hoc internal sensitivity analysis, nested LOOCV was performed using all 34 samples in the primary analysis set and the existing P1 gate. The deterministic cutoff-selection rule used in the nested analysis was formulated after examination of the original dataset to provide a reproducible approximation of the original multi-criteria selection process. This deterministic rule was developed for the post hoc nested analysis and was distinct from the original ChatGPT-assisted recommendation and author decision used to adopt the cutoff of 1000. In each outer iteration, one sample was held out before cutoff selection, and the remaining 33 samples were evaluated at all 65 candidate cutoffs. Eligible cutoffs were required to have a Mann–Whitney U-test P value < 0.05, a likelihood-ratio-test P value < 0.01, a training-set AUC ≥ 0.70, a valid fixed-cutoff inner LOOCV AUC, and a mean retained-event count ≥ 10 across all training samples, with samples containing no retained events counted as zero. Eligible cutoffs were ranked successively by training-set AUC, inner LOOCV AUC, mean retained-event count, and the lower cutoff. If no cutoff met all criteria, the same ranking was applied first among candidates meeting the retained-event requirement and then among candidates with valid AUC estimates. A logistic regression model fitted using evaluable outer-training samples at the selected cutoff was applied once to the held-out sample. A held-out sample with no retained event at the selected cutoff was recorded as unevaluable without imputing a signal intensity of zero. The nested AUC and its 95% CI were calculated from the accumulated evaluable outer predictions using the DeLong method, and the evaluability rate was reported separately. The nominal 95% DeLong CI is reported descriptively and does not explicitly account for dependence among out-of-fold predictions or uncertainty arising from cutoff selection and model fitting.

### 2.9. Assessment of intra- and inter-assay precision

A separate serum sample (Sample 41), which was not included in the clinical analysis, was used to assess intra- and inter-assay precision. The same serum sample was maintained at 4 °C after processing on day 1 and was used for the preparations for day 2. A separate SEC column was used for each replicate preparation.

For each replicate, the procedure from initial SEC preparation through EMARS labeling was performed independently. Samples labeled on day 1 were stored at 4 °C in the dark for approximately 24 h and measured together with the day 2 preparations under the same flow cytometry conditions. P1 and the FITC-H cutoff of 1000 were applied to calculate MFI. Within-day and between-preparation coefficients of variation (CVs) were evaluated descriptively.

### 2.10. Sensitivity analysis of clinical classification

To evaluate the effect of the classification of Samples 5, 92, and 102, which were excluded from the primary analysis, a sensitivity analysis was performed in which these three samples were included in the ILD group. The sensitivity-analysis population therefore comprised 37 samples: 21 ILD and 16 non-ILD samples. P1 and the FITC-H cutoff of 1000 from the primary analysis were applied without modification, and the cutoff was not reselected. MFI values were calculated, and statistical analyses were performed as described for the primary analysis.

### 2.11. Statistical analysis

Final statistical analyses and figure preparation were performed using R version 4.5.3 (R Foundation for Statistical Computing, Vienna, Austria). Receiver operating characteristic curve analysis, AUC calculation, and calculation of 95% CIs using the DeLong method were performed using the pROC package version 1.19.0.1. Continuous variables were compared between two groups using the two-sided Mann–Whitney U test. Profile-likelihood 95% CIs were calculated for odds ratios. Univariable logistic regression models were compared with intercept-only models using the likelihood-ratio test. All tests were two-sided. P values were interpreted descriptively in this exploratory analysis, and no adjustment was made for comparisons across candidate cutoffs or for the cutoff-selection process. The AI-assisted analysis procedure and cutoff-selection method are described in Section 2.8.

## 3. Results

### 3.1. Workflow for proximity labeling and flow cytometric analysis

The overall experimental protocol is illustrated in Fig. 1. To use proximity labeling for EV analysis, EVs were first purified from serum. Because previous studies [14] demonstrated that SEC provides a simple purification method with favorable recovery, SEC was also used in this study. For labeling EVs by proximity labeling followed by flow-cytometric analysis, the workflow comprised the following steps: Step 1, Serum preparation; Step 2, SEC-based EV enrichment; Step 3, the EMARS reaction (for the underlying principle, see Fig. S1) with FT reagent (or, for comparison, conventional labeling with a secondary antibody) followed by removal of excess reagents; and Step 4, flow cytometric analysis using a FACSCanto II flow cytometer. SEC purification was used not only for EV purification from serum but also at each step to remove excess anti-CHL1–HRP (the EMARS probe) and FT, resulting in a total of three SEC procedures within a single workflow (Fig. 1B). For the two excess-reagent removal steps, the SEC resin used for EV purification was washed with PBS and reused.

**Figure 1.**
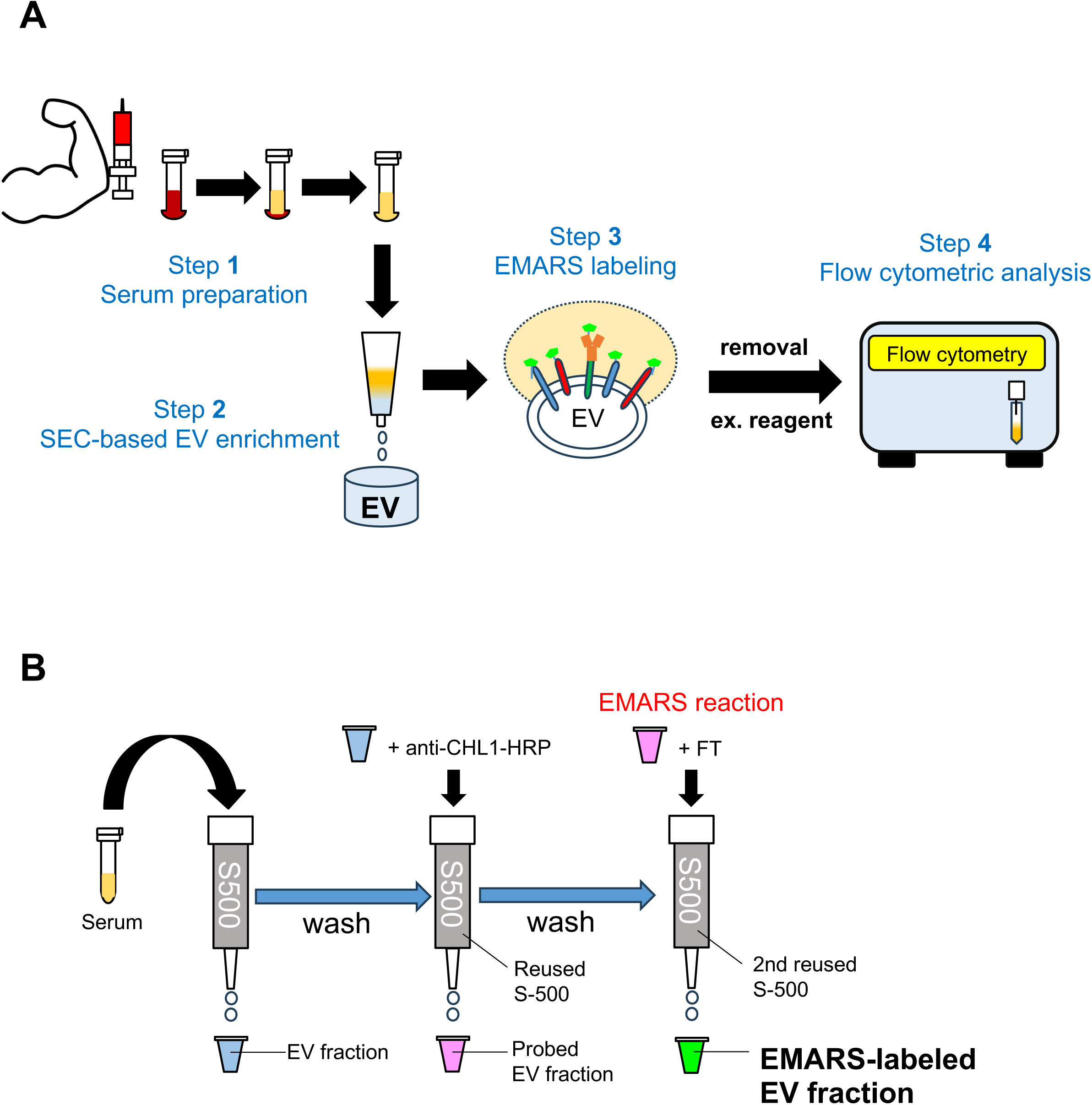
Workflow of EMARS labeling and flow cytometric analysis of serum EV-enriched fractions. (A) Overview of the complete workflow, from sample preparation to flow-cytometric analysis, used in this study. Serum samples were subjected to SEC, followed by CHL1-based EMARS labeling (or conventional anti-CHL1 antibody labeling). In the EMARS reaction, HRP conjugated to the anti-CHL1 antibody activated FT and labeled molecules located near CHL1. Fluorescence was measured by flow cytometry. (B) Schematic representation of the SEC procedure. In this study, Sephacryl S-500 mini-columns were used for three steps: initial recovery of the serum EV-enriched fraction, removal of unbound antibody, and removal of residual EMARS reagents. Rather than using a new column for each step, the column used in the initial step was washed with PBS and reused.

### 3.2. Characterization of serum EVs in the SEC fraction

NTA of a serum EV-enriched fraction prepared by SEC showed a modal diameter of 156.0 nm and a D50 of 240.9 nm, with a heterogeneous distribution of submicron particles (Fig. 2A). TEM revealed vesicle-like particles of various sizes in multiple fields of view (Fig. 2B and Fig. S2). In the Western blot analysis, the EV-positive markers CD63 and integrin alpha 2 were detected, whereas the EV-negative marker beta-actin was not detected (Fig. 2C). These findings indicated the presence of EVs in the recovered serum EV-enriched fractions.

**Figure 2.**
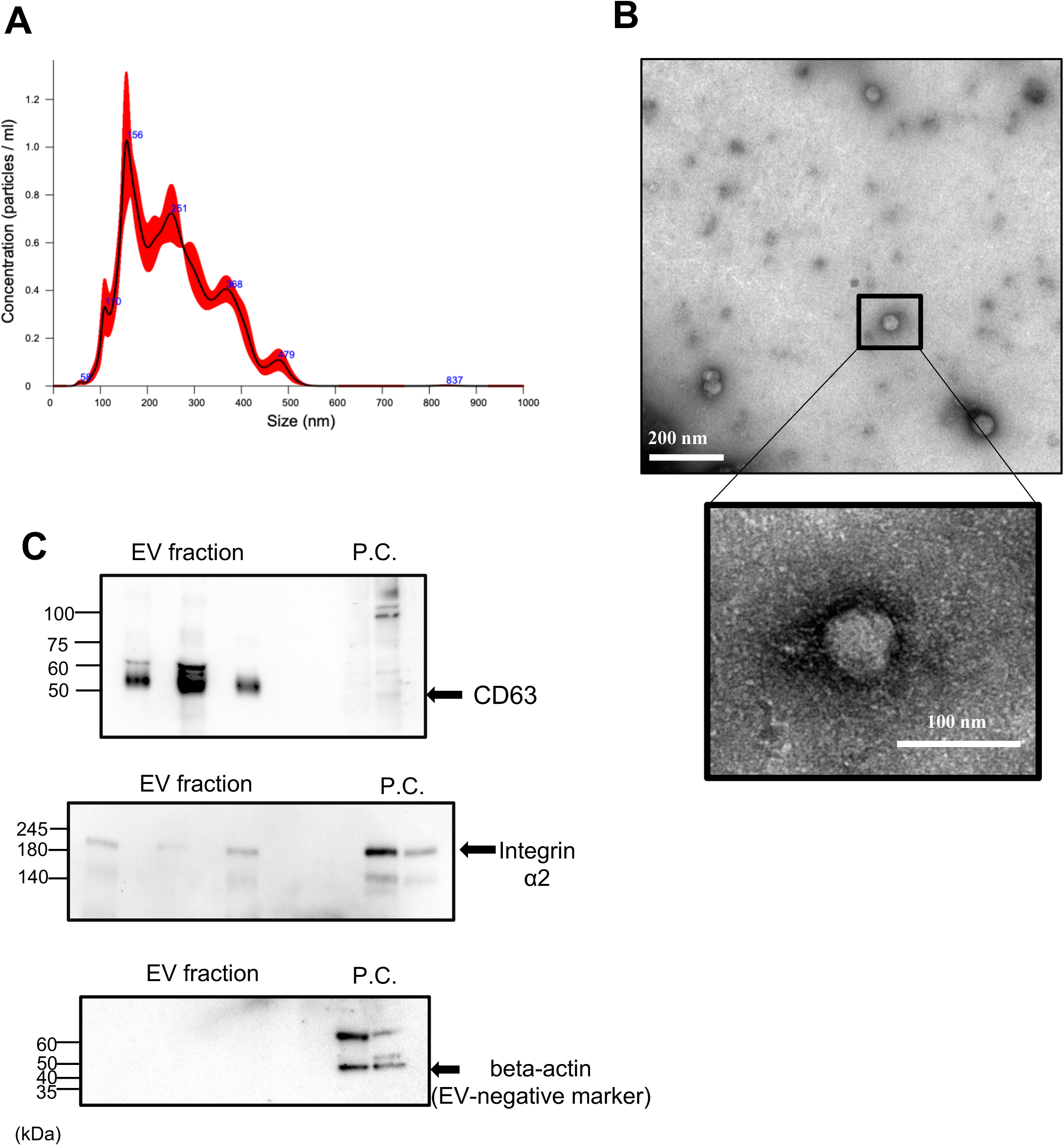
Characterization of SEC-derived serum EV-enriched fractions. (A) Representative particle-size distribution measured by NTA. The values shown in blue indicate the particle size (nm) at the apex of each peak. The red shaded area indicates the dispersion of the particle-size distribution across replicate measurements. (B) Representative TEM images of the serum EV-enriched fraction. The boxed region in the upper image is shown at higher magnification below. Scale bars, 200 nm and 100 nm, respectively. (C) Western blot analysis of CD63 (EV marker), integrin alpha 2 (EV marker), and beta-actin (EV negative marker) in SEC-derived samples and cellular controls. The three EV fraction lanes represent independent SEC fractions 6–8 prepared from serum samples of three different patients. A549 and 293T cell lysates were used as positive controls and are shown from left to right, respectively. Molecular mass markers are indicated in kDa. Uncropped images are shown in Fig. S8. Experiments were performed in duplicate.

### 3.3. Flow cytometric analysis of EMARS-labeled particles in serum EV-enriched fractions

A commercially available mixture of standardized particles (Megamix-Plus SSC particles), previously used for flow-cytometric analysis of EVs, was analyzed using an SSC-H versus FITC-H bivariate scatter plot. Under the manufacturer-recommended instrument settings, the standard particles were successfully resolved and detected (Fig. 3A). Next, using these settings, PBS without EVs was subjected to the EMARS procedure as a negative control experiment and analyzed by flow cytometry (Fig. 3B). Strong background signals, presumably originating from S-500 resin and/or reagents, were detected in the low-FITC-H/low-SSC-H and high-SSC-H/low-FITC-H regions. Using the reference-particle and PBS control measurements, P1 was positioned to encompass the 0.16-, 0.20-, and 0.24-µm reference-particle distributions while excluding the prominent low-FITC-H/low-SSC-H and high-SSC-H/low-FITC-H background regions. Events within P1 were subsequently analyzed. Next, serum EV-enriched fractions were prepared and labeled by EMARS according to the workflow described above, and the resulting samples were analyzed using the same instrument settings. In the P1 gate, particles with high FITC-H signals that were absent from the negative control were detected (Fig. 3C).

**Figure 3.**
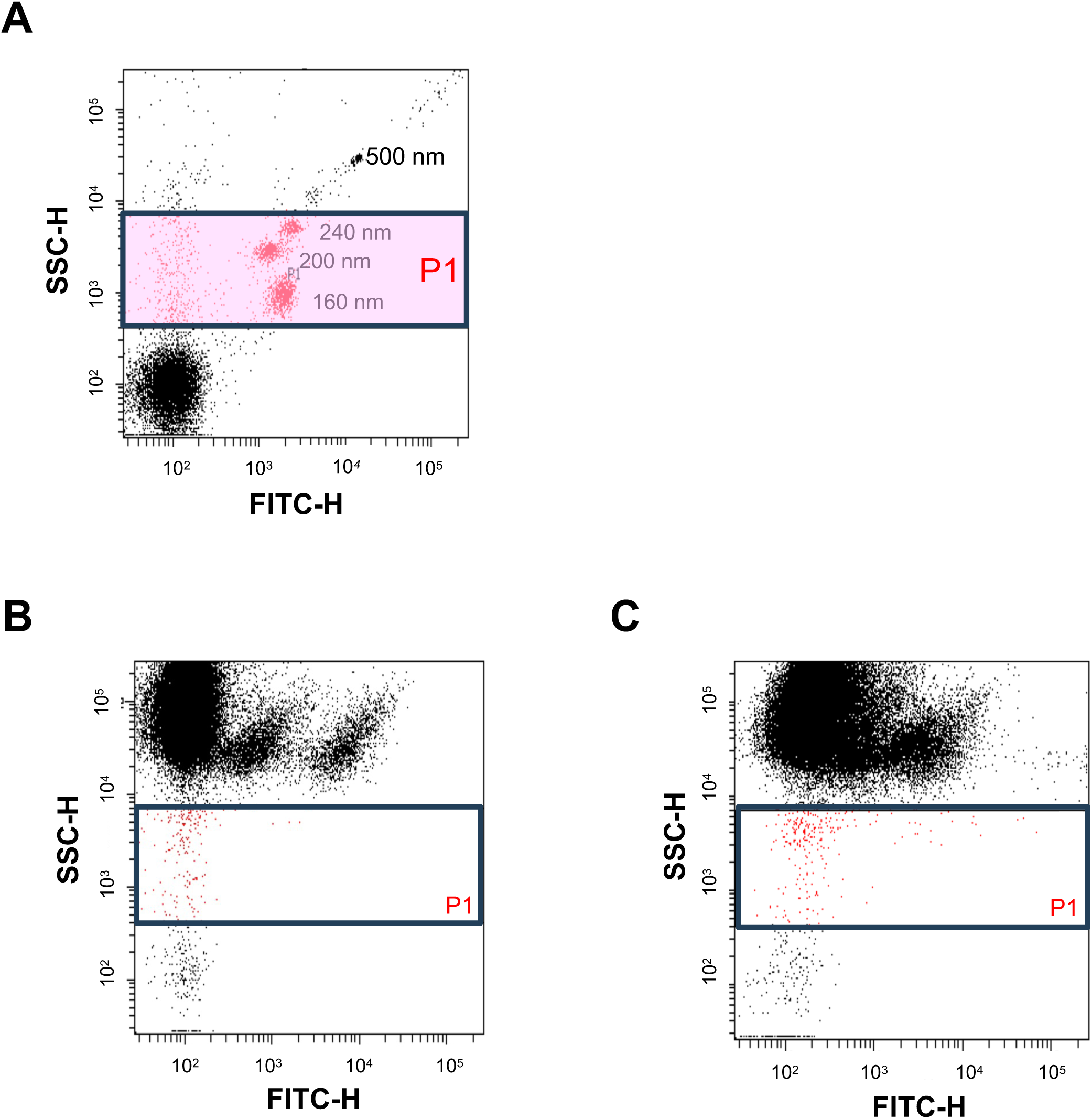
Flow cytometric analysis of serum EV-enriched fractions following EMARS labeling. (A) SSC-H versus FITC-H plot of Megamix-Plus SSC reference particles. The light-red area denotes the operational P1 gate, positioned with reference to the 0.16-, 0.20-, and 0.24-µm SSC reference-particle distributions and the PBS process control. (B) SSC-H versus FITC-H plot of the PBS control (sample without EVs). The lower boundary of P1 was set to include the distribution of 0.16-µm particles while excluding low-FITC-H/low-SSC-H events. The upper boundary excluded the high-SSC-H/low-FITC-H region that was prominent in the PBS control. (C) SSC-H versus FITC-H plot of a representative serum EV-enriched fraction after EMARS labeling. A session-specific P1 gate was defined using the reference particles and PBS control and subsequently applied to serum samples acquired during the same analytic session.

### 3.4. Comparison of labeling efficacy between the conventional method and EMARS

PBS controls and serum EV-enriched fractions were analyzed after conventional antibody labeling or CHL1-targeted EMARS labeling (Fig. 4A). With conventional labeling, the MFI was 910 for the PBS control and 1099 for the serum EV-enriched fraction, corresponding to a serum-to-PBS ratio of 1.21. After EMARS labeling, the corresponding values were 356 and 2234, yielding a serum-to-PBS ratio of 6.28 (Fig. 4B). Numerically, fluorescence in the serum EV-enriched fraction was 2.03-fold higher and fluorescence in the PBS control was 2.56-fold lower after EMARS labeling than after conventional labeling. Because the fluorophores and labeling reactions differed, absolute fluorescence intensities were not compared directly between the two methods.

**Figure 4.**
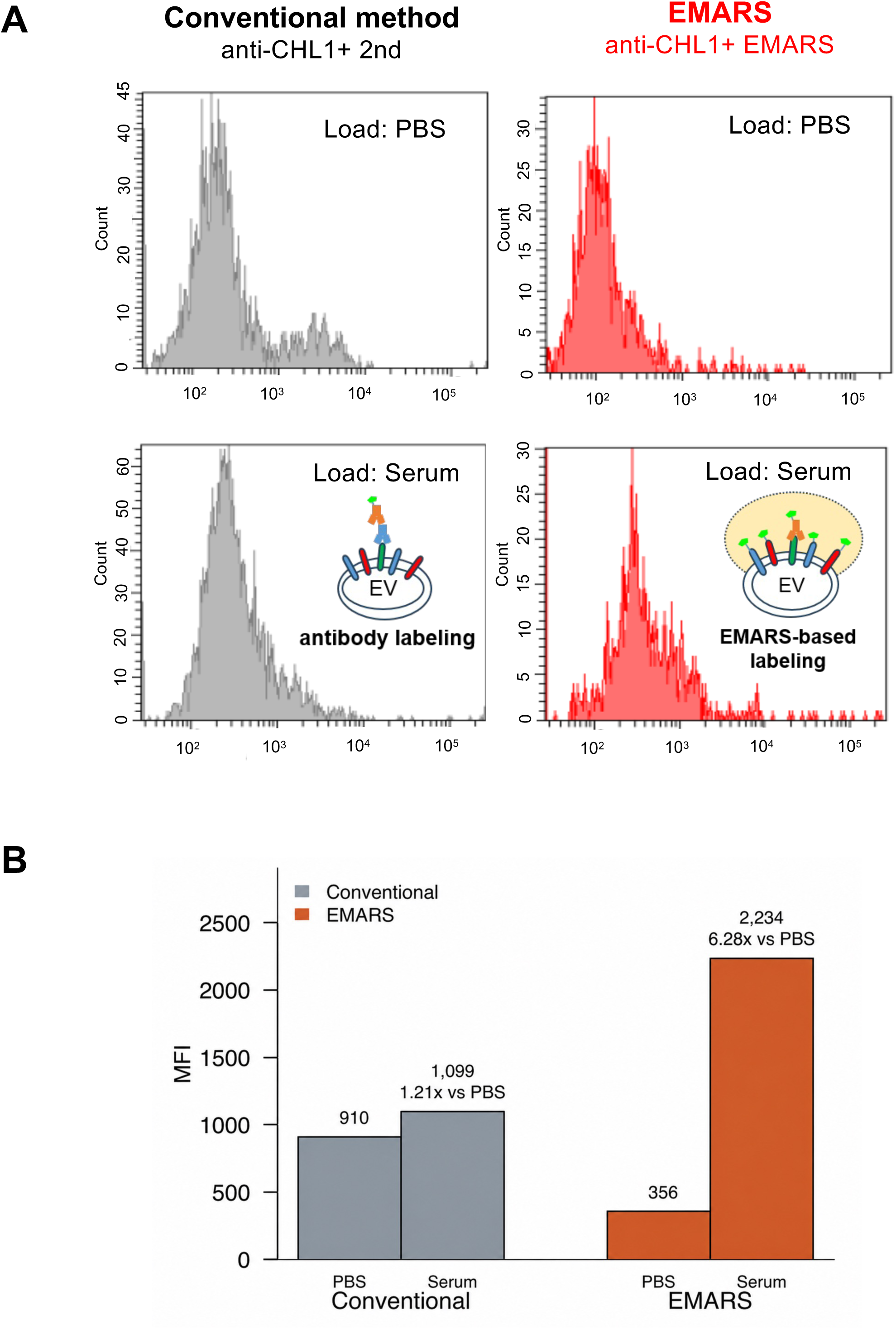
Comparison of fluorescence ratios between conventional and EMARS labeling. (A) Representative fluorescence histograms for PBS control and serum EV-enriched fraction after conventional anti-CHL1 antibody labeling (left) or EMARS labeling (right). PBS controls are shown in the upper panels and serum-derived samples in the lower panels. Experiments were performed in duplicate. (B) Fluorescence intensities from the representative measurements shown in (A). The serum-to-PBS ratio was 1.21 for conventional labeling and 6.28 for EMARS labeling. Because the methods used different fluorescence-labeling reactions, absolute fluorescence intensities were not compared directly.

### 3.5. AI-assisted selection of the FITC-H cutoff

After establishing the SSC-H P1 gate, serum EV-enriched fractions from patients with pulmonary disease were labeled by EMARS and analyzed by flow cytometry. Using all events within P1 before application of a FITC-H cutoff, the cancer versus non-cancer comparison yielded a Mann–Whitney U-test P value of 0.320 and a logistic-regression likelihood-ratio-test P value of 0.197 (Fig. S3A). For the ILD versus non-ILD comparison, the corresponding P values were 0.116 and 0.0432, respectively (Fig. S3B). Because the initial cancer comparison showed little separation, whereas the ILD comparison differed according to the statistical method, the subsequent exploratory cutoff analysis focused on ILD status.

Incrementally testing candidate FITC-H thresholds and calculating the statistical and operational measures for each threshold was labor-intensive. We therefore used the AI-assisted cutoff-selection workflow shown in Fig. 5A. In the full-dataset analysis, ChatGPT compared 65 candidate FITC-H values and recommended FITC-H = 1000 based on the balance among ILD versus non-ILD discrimination, fixed-cutoff cross-validated performance, and event retention. The authors reviewed the complete cutoff-specific results and adopted this value as the exploratory, data-selected cutoff. At FITC-H = 1000, the AUC obtained from the full-dataset analysis was 0.846, and the fixed-cutoff LOOCV AUC was 0.764 (Fig. 5B). The event distribution within P1 and the selected cutoff are shown in the right panel of Fig. 5B. Because cutoff selection was not repeated within the leave-one-out iterations, the fixed-cutoff LOOCV does not estimate the performance of the complete cutoff-selection procedure. Increasing the cutoff reduced the number of events retained for analysis and increased the proportion of samples with no retained events (Figs. 5C and S4A). Before application of the cutoff, the median number of events in P1 was 45 (range, 14–295) among all 34 samples. At FITC-H = 1000, 28 evaluable samples retained a median of 9.5 events (range, 1–39). Six samples, comprising three ILD and three non-ILD samples, had no retained events and were therefore not evaluable for MFI at this cutoff. The median total P1 event count was lower in these samples than in the evaluable samples (31.5 [range, 14–97] versus 62 [range, 15–295]), although the ranges overlapped. To assess intra- and inter-assay precision, serum from Sample 41 was analyzed in duplicate on the same day using identical sample-preparation and flow-cytometric procedures. The assay was then repeated on a separate day using the same serum sample under identical experimental conditions. The duplicate MFI values were 2900 and 3169 on Day 1 (within-day CV, 6.3%) and 2516 and 3085 on Day 2 (within-day CV, 14.4%). The between-day CVs calculated for the first and second replicate measurements were 10.0% and 1.9%, respectively (Fig. S4B).

**Figure 5.**
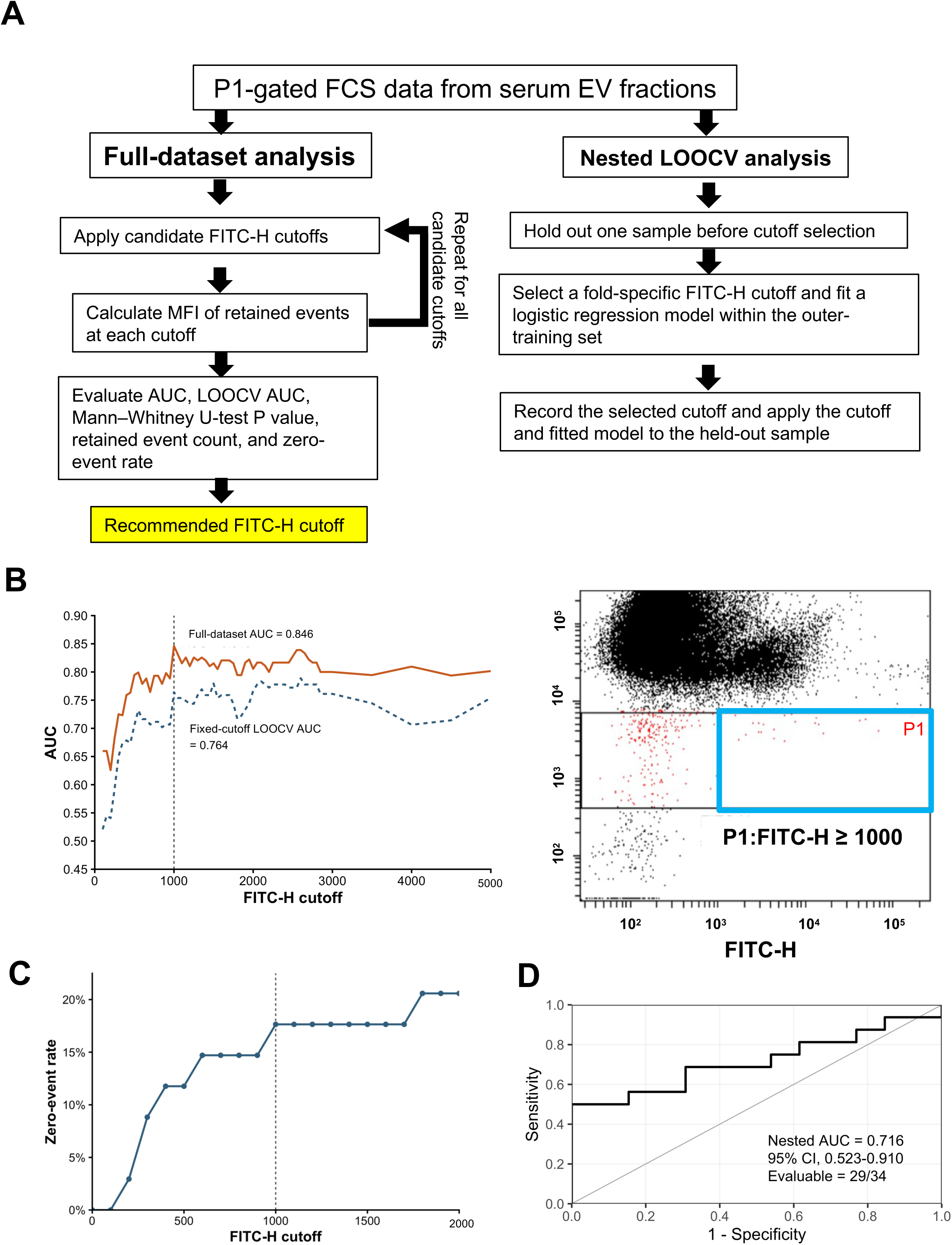
Full-dataset and nested analyses for AI-assisted FITC-H cutoff selection. (A) Workflow of the AI-assisted cutoff-selection analysis. In the full-dataset analysis (left), all 34 samples entered the cutoff-comparison procedure, and cutoff-specific statistics were calculated using the evaluable samples at each cutoff. ChatGPT returned a recommended cutoff, and the authors reviewed the complete results before adopting FITC-H = 1000. The nested LOOCV (right) was performed separately as a post hoc internal sensitivity analysis after the cutoff of 1000 had been adopted and did not contribute to its initial selection. (B) Full-dataset AUC and fixed-cutoff LOOCV AUC across candidate cutoffs (left). At FITC-H = 1000, the AUC obtained from the full-dataset analysis was 0.846 and the fixed-cutoff LOOCV AUC was 0.764. The AUC was not adjusted for cutoff selection. Because cutoff selection was not repeated within each leave-one-out iteration, the fixed-cutoff LOOCV result does not provide an unbiased estimate of the performance of the complete selection procedure. The event distribution within P1 and the selected cutoff area (blue square) at FITC-H = 1000 are shown on the right. (C) Proportion of samples with no events at or above each cutoff, shown over the range FITC-H 0–2000. Such samples were excluded from the analysis at that cutoff. (D) ROC curve generated from the 29 evaluable outer hold-out predictions in the post hoc nested LOOCV sensitivity analysis. The nested AUC was 0.716 (95% CI, 0.523–0.910); five of 34 hold-out samples had no retained events at the fold-specific selected cutoff and were not evaluable.

In the separate post hoc nested LOOCV sensitivity analysis, cutoff selection was repeated within each outer fold. Twenty-nine of 34 outer hold-out samples were evaluable (85.3%). The nested AUC was 0.716 (95% CI, 0.523–0.910; Fig. 5D). The median selected cutoff was 1000 (range, 550–1000), and cutoff 1000 was selected in 31 of 34 folds (Fig. S5). Thus, cutoff 1000 was the most frequently selected value, whereas the nested AUC was lower than the AUC obtained from the full-dataset analysis.

We additionally explored the contribution of the P1 gate by comparing analyses performed with and without P1 at the previously selected exploratory cutoff of FITC-H = 1000. Across the evaluated cutoff range, the AUC calculated in matched evaluable samples was higher with P1 than without P1 (Fig. S6). The same 28 samples were analyzed with and without P1 at FITC-H = 1000. With P1, the AUC was 0.846 and the fixed-cutoff LOOCV AUC was 0.764; without P1, the corresponding values were 0.605 and 0.487. In this dataset, discrimination was higher when the P1 gate was applied. Because the cutoff was selected within the P1-based workflow, this comparison is exploratory and does not establish the general superiority of P1 gating.

### 3.6. Exploratory association between CHL1-targeted particle fluorescence and ILD

After applying P1 and FITC-H = 1000, the MFI of retained events in serum EV-enriched fractions was compared between the ILD and non-ILD groups. The primary analysis included 34 samples: 18 ILD and 16 non-ILD samples, the latter comprising 11 thoracic malignancies and five other respiratory diseases (Table 1). Twenty-eight samples were evaluable, including 15 ILD and 13 non-ILD samples. MFI was higher in the ILD group (Mann–Whitney U test, P = 0.002; Fig. 6A). The AUC obtained from the full-dataset analysis, calculated using these same 28 samples, was 0.846 (95% CI, 0.704–0.988; Fig. 6B), and the fixed-cutoff LOOCV AUC was 0.764. Because FITC-H = 1000 was selected using the same dataset, the P value and AUC were not adjusted for the cutoff-selection process. In univariable logistic regression, the odds ratio per 100-unit increase in MFI was 1.031 (95% profile-likelihood CI, 1.010–1.067; likelihood-ratio test, P = 0.000543; Fig. 6C). Among the 28 evaluable samples, nine were from patients with cancer and 19 from patients without cancer. All nine cancer samples were in the non-ILD group, whereas 15 of the 19 non-cancer samples were in the ILD group. Median MFI was 3716.0 (range, 1432.5–9911.2) in the cancer samples and 6938.9 (range, 1959.4–31919.2) in the non-cancer samples (Fig. S7A). Within the non-ILD group, median MFI was 3716.0 (range, 1432.5–9911.2) in the nine cancer samples and 3580.3 (range, 1959.4–6193.5) in the four samples from patients with other respiratory diseases.

**Figure 6.**
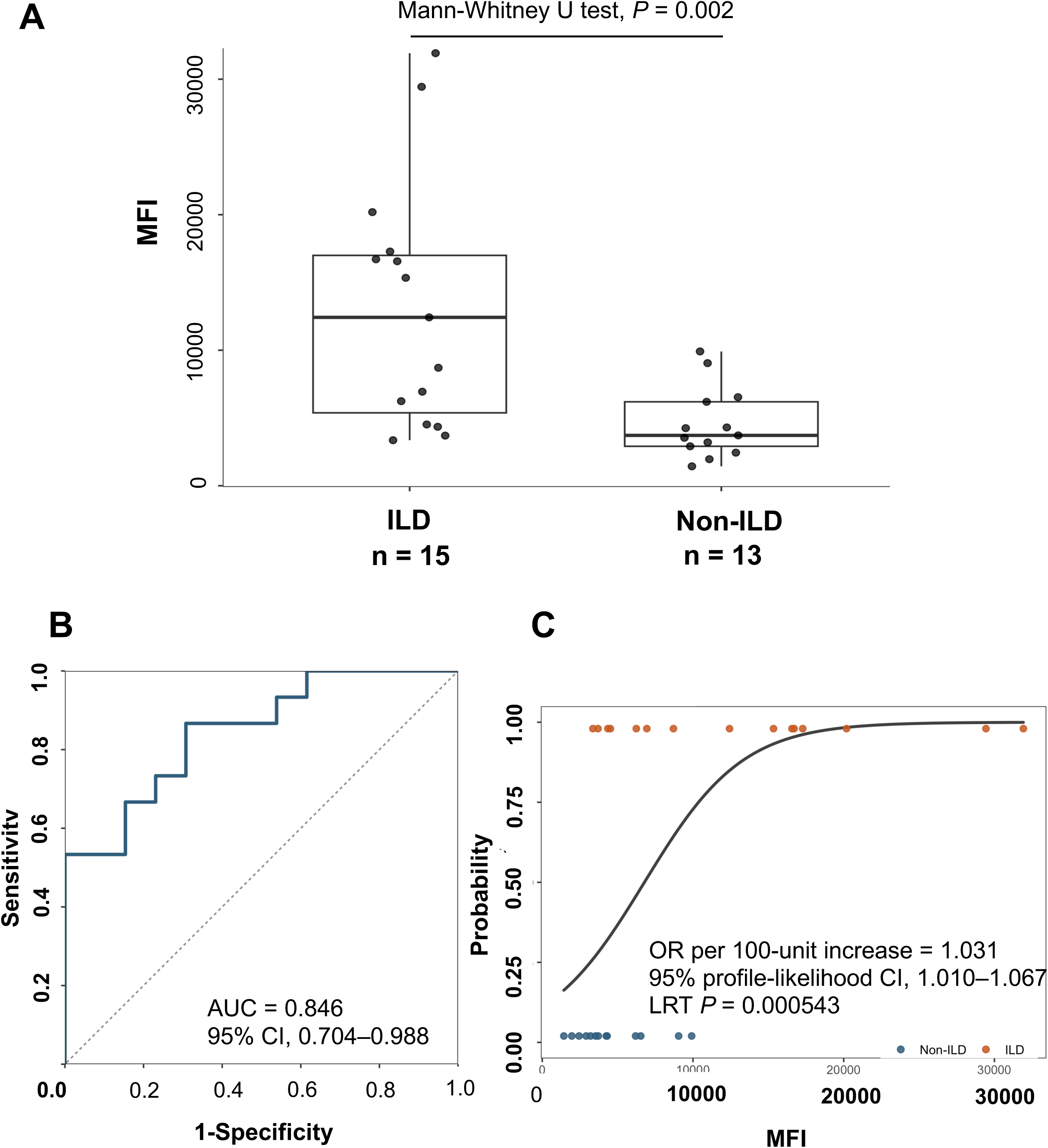
Exploratory comparison of CHL1-targeted particle fluorescence between ILD and non-ILD samples. (A) MFI of retained events in serum EV-enriched fractions from the ILD (n = 15) and non-ILD (n = 13) groups after application of P1 and FITC-H = 1000. Of the 34 samples in the primary analysis set, 28 contained at least one P1 event with FITC-H ≥ 1000 and were included in the MFI comparison. Six samples, comprising three ILD and three non-ILD samples, had no retained events and were not evaluable for MFI. Groups were compared using the Mann–Whitney U test (P = 0.002). (B) ROC curve for discrimination of ILD from non-ILD. The AUC obtained from the full-dataset analysis was 0.846 (95% CI, 0.704–0.988). The P value in panel A and the AUC in panel B were not adjusted for cutoff selection. (C) Univariable logistic regression of ILD status on MFI. The odds ratio was 1.031 per 100-unit increase (95% profile-likelihood CI, 1.010–1.067; likelihood-ratio-test P = 0.000543). Blue dots indicate non-ILD samples, and orange dots indicate ILD samples.

When the three samples excluded from the primary analysis were added to the ILD group, 31 samples were evaluable, including 18 ILD and 13 non-ILD samples. MFI remained higher in the ILD group (Mann–Whitney U test, P = 0.000535; Fig. S7B). The AUC was 0.872 (95% CI, 0.751–0.993), and the fixed-cutoff LOOCV AUC was 0.803.

## 4. Discussion

The present study established a workflow for measuring target-associated fluorescence from serum EV-enriched fractions using proximity labeling (EMARS) and a conventional flow cytometer. The workflow integrates preparation of serum EV-enriched fractions by SEC, enhanced fluorescein labeling by EMARS, event selection using P1, and AI-assisted selection of a FITC-H cutoff. The principal contribution is the use of EMARS to enhance target-associated fluorescence to a level that could be recorded as event-level FCS data on an existing conventional flow cytometry system; P1 and the AI-assisted cutoff then determine which events enter the analysis. EMARS enhances fluorescence intensity, whereas P1 (Fig. 3B) and the FITC-H cutoff exclude unwanted events originating from debris and other non-target particles. The cutoff was selected while explicitly considering the reduction in retained events and evaluable samples at higher thresholds. At the recommended cutoff (FITC-H = 1000), the full-dataset analysis yielded evaluable MFI values in 28 of 34 clinical samples and an AUC of 0.846. In the post hoc nested LOOCV sensitivity analysis, 29 of 34 outer hold-out samples were evaluable, the AUC was 0.716 (95% CI, 0.523–0.910), and cutoff 1000 was selected in 31 of 34 folds. The lower nested AUC indicates more modest clinical discrimination when cutoff selection is incorporated into internal validation, while the repeated selection of cutoff 1000 supports its stability within this dataset.

SEC is well suited to the preparation of serum EV-enriched fractions from limited volumes, but its performance depends on the column and the biological matrix. Karimi et al. showed overlap between EV- and lipoprotein-associated components after SEC and improved their separation by combining SEC with density-based fractionation [17]. A more recent comparison confirmed the practical accessibility and EV yield of SEC from 1 mL of plasma while also detecting residual apolipoproteins in the SEC products [18]. Thus, the persistence of lipoprotein overlap is a general limitation of SEC rather than a problem unique to the present column. In this workflow, the three SEC steps served distinct purposes: initial recovery of a serum EV-enriched fraction, removal of unbound antibody, and removal of residual EMARS reagents. To ensure experimental reproducibility, appropriate next steps include standardizing column packing, monitoring flow rates and recovery, and evaluating prepacked columns.

Weak fluorescence in fluorescence-labeled EVs has been a central constraint in conventional EV flow cytometry [4,6]. In the present reaction, HRP conjugated to the anti-CHL1 antibody activates FT and deposits multiple fluorescein molecules near the antibody-bound target (Fig. S1). The resulting fluorescence can be enhanced and read directly on the FITC channel. In the representative comparison, the detection ratio increased approximately fivefold after EMARS labeling compared with conventional antibody labeling. Tyramide amplification has similarly increased fluorescence in other single-particle EV analyses [19]. By combining this signal-amplification approach with an AI-assisted cutoff for flow-cytometric analysis, this study provides a basis for the simple analysis of clinical specimens and for the future development of EV-based diagnostics.

The EMARS fluorescein signal contains information that differs from a direct measurement of CHL1 abundance. Tyramide-based EMARS labels proteins located near an HRP-conjugated antibody, and the measured fluorescence can consequently reflect the presence and distribution of CHL1-bearing EVs together with the local molecular environment surrounding CHL1. In the previous EV study, the abundance of products labeled by CHL1-initiated EMARS did not necessarily parallel the abundance of the corresponding proteins in the total serum EV-enriched fractions [14]. This property is relevant to the present method because an EV may express several proteins within the effective labeling range of the CHL1-bound antibody, allowing their combined fluorescein deposition to contribute to particle-associated fluorescence. Because the methodological objective was to evaluate the strength of fluorescence associated with the retained events, rather than event number, their MFI was used as the sample-level readout of EV fluorescence.

CHL1 was selected as a model target because it had previously been used to examine lung cancer-associated EVs [14]. That study specifically found higher levels of serum EVs coexpressing CHL1 and caspase-14 in patients with lung cancer than in healthy individuals [14]. In contrast, when focusing solely on CHL1 expression in serum-derived EVs from patients with lung cancer, no significant difference has been reported compared with healthy individuals [14]. In the present exploratory analysis, CHL1 served as the initiating target for EMARS labeling, and the observed association therefore concerns CHL1-initiated EMARS fluorescence. Given the labeling principle of EMARS, this signal may reflect not only CHL1 abundance but also disease-associated changes in the local molecular environment surrounding CHL1. Changes in circulating EV proteins [20,21] and airway EV surface-marker profiles [22] have been reported in ILD, providing a biological basis for investigating EV-associated molecular changes in these diseases. The present result identifies an exploratory signal for further study and is not intended to validate CHL1-initiated EMARS fluorescence as an ILD diagnostic biomarker. The observed association with ILD requires validation in independent cohorts, together with further assessment of the identity of retained events and of assay specificity. Future studies using additional disease-relevant surface targets, alone or in combination with CHL1, may clarify whether other targets, or the molecular environments surrounding them, provide more informative signals.

Several sensitive EV assays address complementary analytical needs. ExoPLA permits event-based detection of defined surface-marker combinations, ExoScreen provides a rapid proximity assay using small volumes of unpurified serum, and SAViA supports sensitive plate-based profiling [23–25]. A practical feature of the present workflow is that EMARS-deposited fluorescein is acquired directly using the FITC channel of a conventional flow cytometer. Although an HRP-conjugated antibody must be prepared and validated for each target, the downstream labeling and acquisition workflow can be retained when the surface target is changed. Flow cytometry also preserves the scatter and fluorescence values of recorded events in the FCS file. The ungated files retained in this study can therefore be reanalyzed using alternative gates, cutoffs, and event-distribution features within the parameters originally acquired. Moreover, the stored ungated FCS files permit future AI-assisted multidimensional gating to examine whether events in the regions currently treated as background carry target-associated fluorescence, extending event selection from a manually defined two-parameter region to a higher-dimensional one. Self-organizing maps, automated EV fluorescence gating, and machine-learning analysis of multiparameter event distributions illustrate possible directions for this development [26–28].

This study also has limitations that define the next stage of analytical development. No serial dilution series was performed to assess swarm detection. The P1-gated, FITC-H ≥ 1000 population was sparse after SEC purification and EMARS labeling; further dilution would have yielded too few retained events to calculate robust sample-level MFI values. Thus, dilution-based assessment of coincidence or swarm detection was not feasible under the present assay conditions. Accordingly, swarm detection and contributions from particle aggregates cannot be excluded, and future assay development should aim to increase target-event recovery, for example through optimization of EV enrichment, labeling efficiency, background reduction, and acquisition duration or volume. These improvements would enable serial dilution testing, detergent-sensitivity controls, and more robust estimation of both event concentration and fluorescence-intensity distributions. The clinical analysis was small, single-center, and exploratory. Post hoc nested LOOCV yielded an AUC of 0.716 with a wide 95% CI and five unevaluable outer hold-out samples; moreover, the deterministic selection rule was formulated after examination of the original dataset. Thus, the nested analysis is an internal sensitivity analysis rather than independent validation, and the present results are insufficient to support diagnostic use. At the fixed cutoff of 1000, six of 34 samples were not evaluable for MFI. NTA, TEM, and western blotting were consistent with the presence of EVs in the SEC fraction but did not establish the identity of every flow-cytometric event, and co-isolated lipoproteins, protein aggregates, EV aggregates, and coincident events cannot be excluded. In particular, EV aggregates may entrap CHL1-expressing EVs intended for analysis, potentially resulting in their loss.

Although the present clinical results are insufficient to support diagnostic use, this study establishes a new proximity-labeling workflow for recording target-associated fluorescence from serum EV-enriched fractions using widely available flow cytometers. Further analytical validation, including target-specific controls, assessment of swarm detection, standardized instrument calibration, and independent clinical validation, will be required to define its future diagnostic applicability.

## CRediT authorship contribution statement

L. K.: Conceptualization, Methodology, Validation, Formal analysis, Investigation, Data curation, Visualization, Writing - original draft, Project administration. Ke. I.: Methodology, Investigation, Writing - review & editing. M. Ku.: Investigation, Writing - review & editing. R. Sa.: Investigation, Resources, Writing - review & editing. S. S., M. S., H. I., R. Se., I. H., and H. N.: Investigation, Resources, Data curation. M. Ka.: Methodology, Formal analysis, Writing - review & editing. K. K.: Investigation, Visualization, Writing - review & editing. Y. M.: Methodology, Resources, Supervision, Writing - review & editing. M. N.: Supervision, Writing - review & editing. T. S.: Supervision, Writing - review & editing. Ko. I.: Funding acquisition, Resources, Writing - review & editing. Y. K.: Methodology, Formal analysis, Supervision, Writing - review & editing. K. N.: Conceptualization, Resources, Supervision, Writing - review & editing. N. K.: Conceptualization, Methodology, Investigation, Validation, Formal analysis, Resources, Writing - review & editing, Supervision, Project administration.

## Data availability

De-identified data are available from the corresponding author upon reasonable request, subject to institutional ethics and data-sharing requirements.

## Acknowledgments

We thank the Saitama Medical University Biomedical Research Center for providing general technical assistance. We also thank Egret-Lab Co., Ltd. for performing nanoparticle tracking analysis as a contracted service.

## Declaration of generative AI and AI-assisted technologies in the manuscript preparation process

During the preparation of this work, the authors primarily used ChatGPT (OpenAI) to assist with computational analysis, organization of analytical outputs, literature checking, and language drafting and editing. OpenAI Codex (OpenAI), Claude (Anthropic), and Perplexity (Perplexity AI) were also used to assist with document organization, literature checking, and language review. The analytical use of ChatGPT is described separately in the Materials and methods. The statistical processing was checked and the statistical results were verified by the authors using R. After using these tools and services, the authors reviewed and verified the inputs, code, outputs, references, interpretations, and manuscript content as needed and take full responsibility for the content of the published article.

## Abbreviations

AI: artificial intelligence
AUC: area under the receiver operating characteristic curve
CHL1: close homolog of L1
CI: confidence interval
CV: coefficient of variation
EMARS: enzyme- mediated activation of radical sources
EV: extracellular vesicle
FCS: flow cytometry standard
FT: fluorescein-tyramide
HRP: horseradish peroxidase
ILD: interstitial lung disease
LOOCV: leave-one-out cross-validation
LRT: likelihood-ratio test
MFI: mean fluorescence intensity
NTA: nanoparticle tracking analysis
PBS: phosphate-buffered saline
SEC: size-exclusion chromatography
TBST: Tris-buffered saline containing Tween 20
TEM: transmission electron microscopy.

