## Supplementary Figs for "Proximity Labeling-Assisted Flow Cytometry Improves Fluorescence Contrast in Extracellular Vesicle-Enriched Fractions"

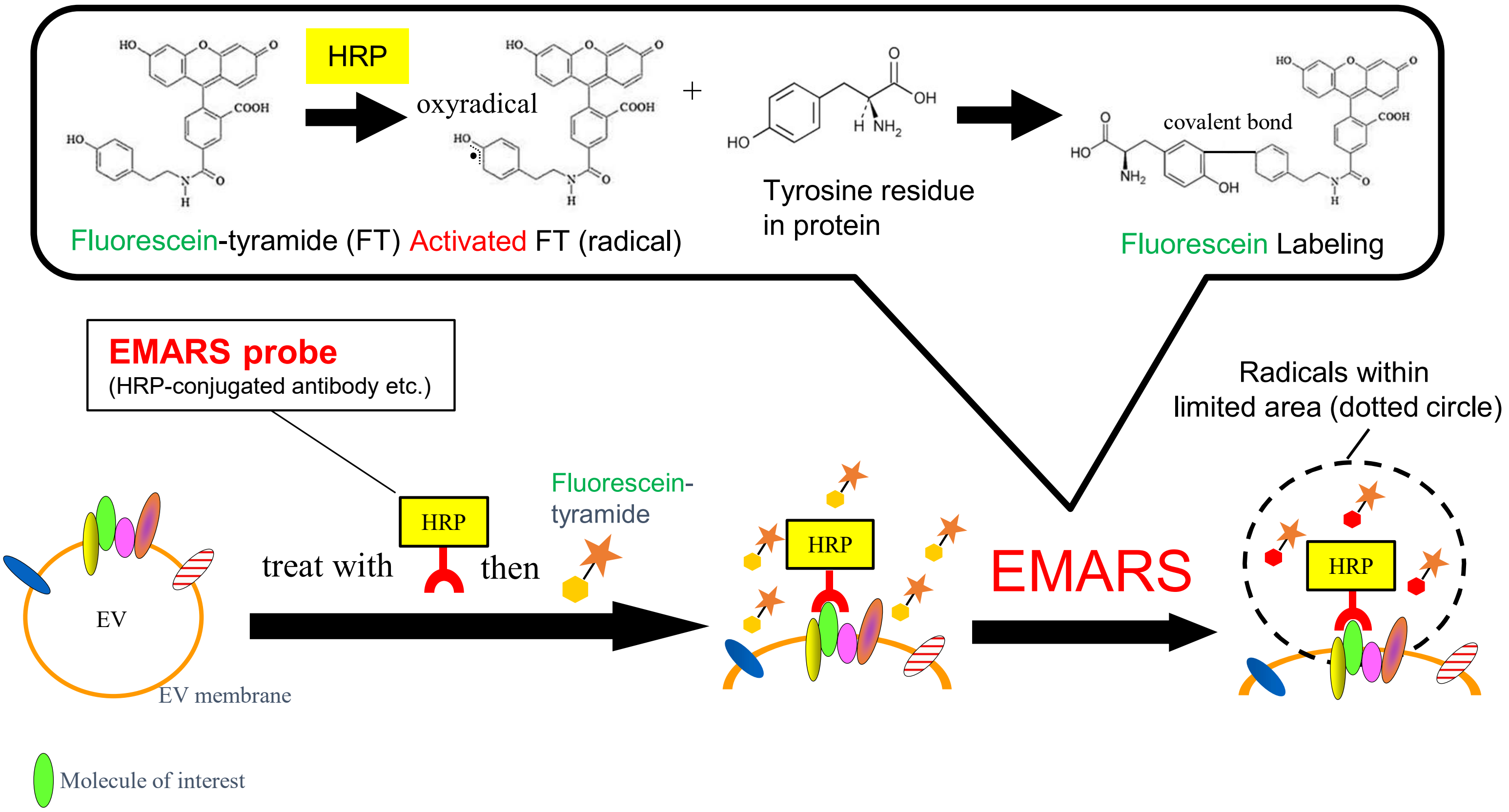


**Figure S1. Principle of EMARS labeling**

An HRP-conjugated antibody binds to a molecule of interest on the cell or nanoparticle surface. In the presence of hydrogen peroxide, HRP activates FT to generate short-lived radicals, which covalently attach fluorescein to molecules located near the antibody. The diagram illustrates the proposed basis for increasing fluorescein labeling around the target molecule.

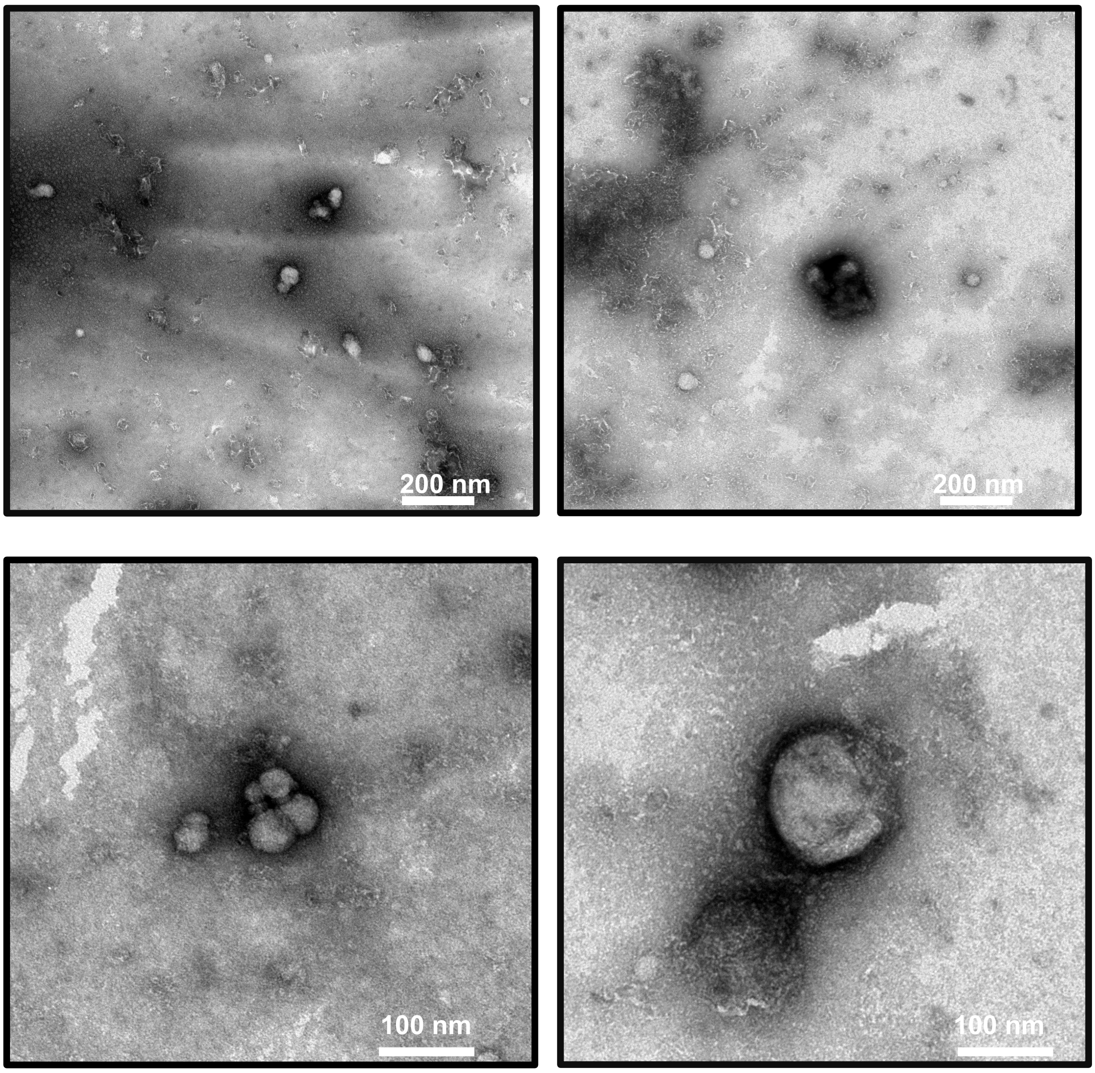


**Figure S2. Additional TEM images of the serum EV-enriched fractions**
Additional TEM images for Fig. 2B show vesicle-like particles in the SEC-derived serum fraction. Scale bars are indicated in the images. White bar: 200 nm or 100 nm.

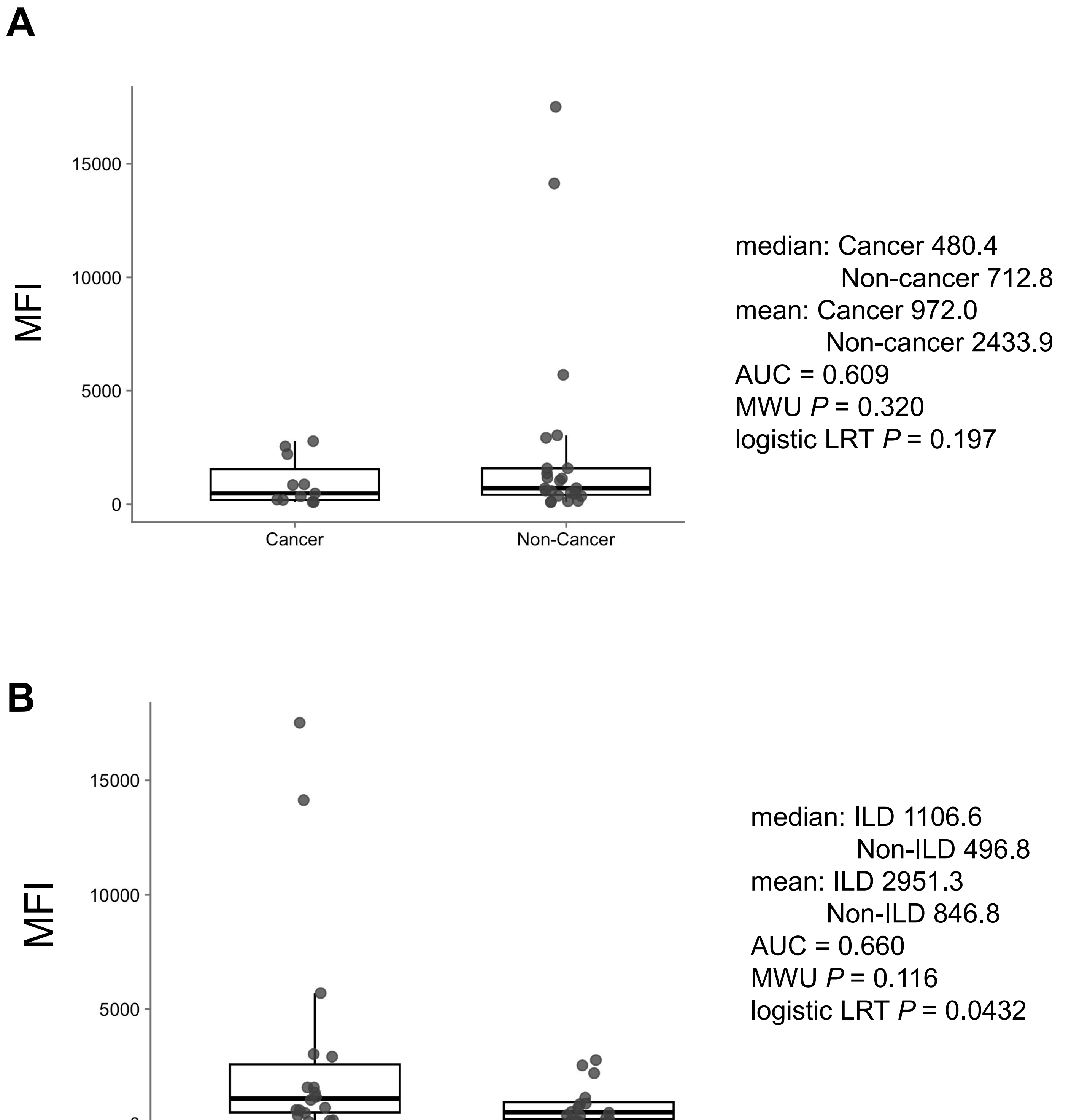


**Figure S3. Comparison of MFI in serum EV-enriched fractions without AI-assisted cutoff selection**

(A, B) Comparisons of MFI in serum EV-enriched fractions between cancer and non-cancer samples (A) and between ILD and non-ILD samples (B). For the cancer versus non-cancer comparison, the P values from the Mann–Whitney U (MWU) test and the likelihood-ratio test for logistic regression were 0.320 and 0.197, respectively. For the ILD versus non-ILD comparison, the corresponding P values were 0.116 and 0.0432, respectively.

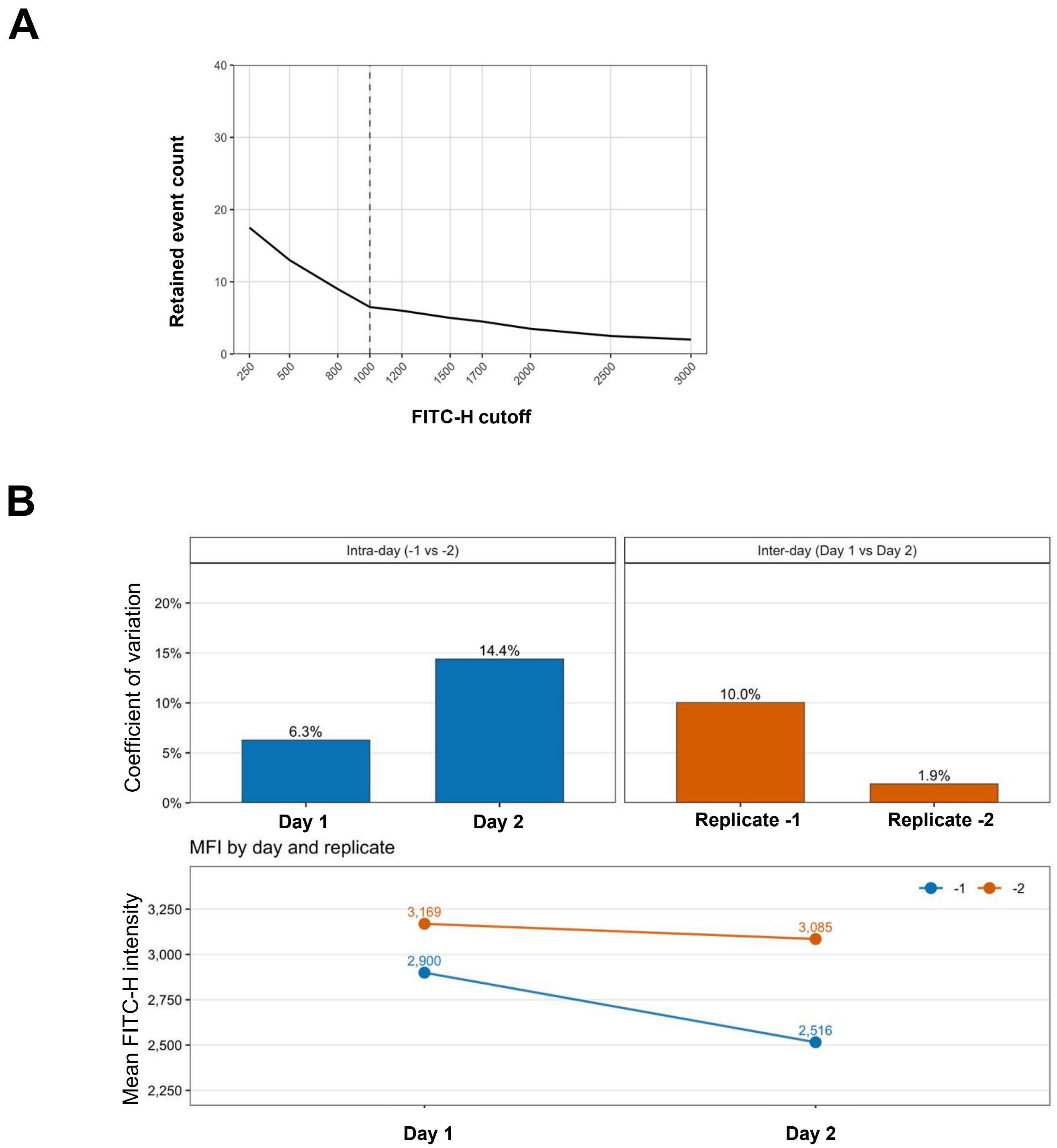


**Figure S4. AI-assisted flow-cytometric analysis under different experimental conditions**

(A) Number of events retained across cutoffs. (B) Assessment of intra- and inter-assay precision. Duplicate SEC columns were prepared and processed on day 1 or day 2. Samples from both days were analyzed in the same flow-cytometry run. Sample 41 showed within-day CVs of 6.3% and 14.4% on days 1 and 2, respectively, and between-day CVs of 10.0% and 1.9% for the corresponding replicates.

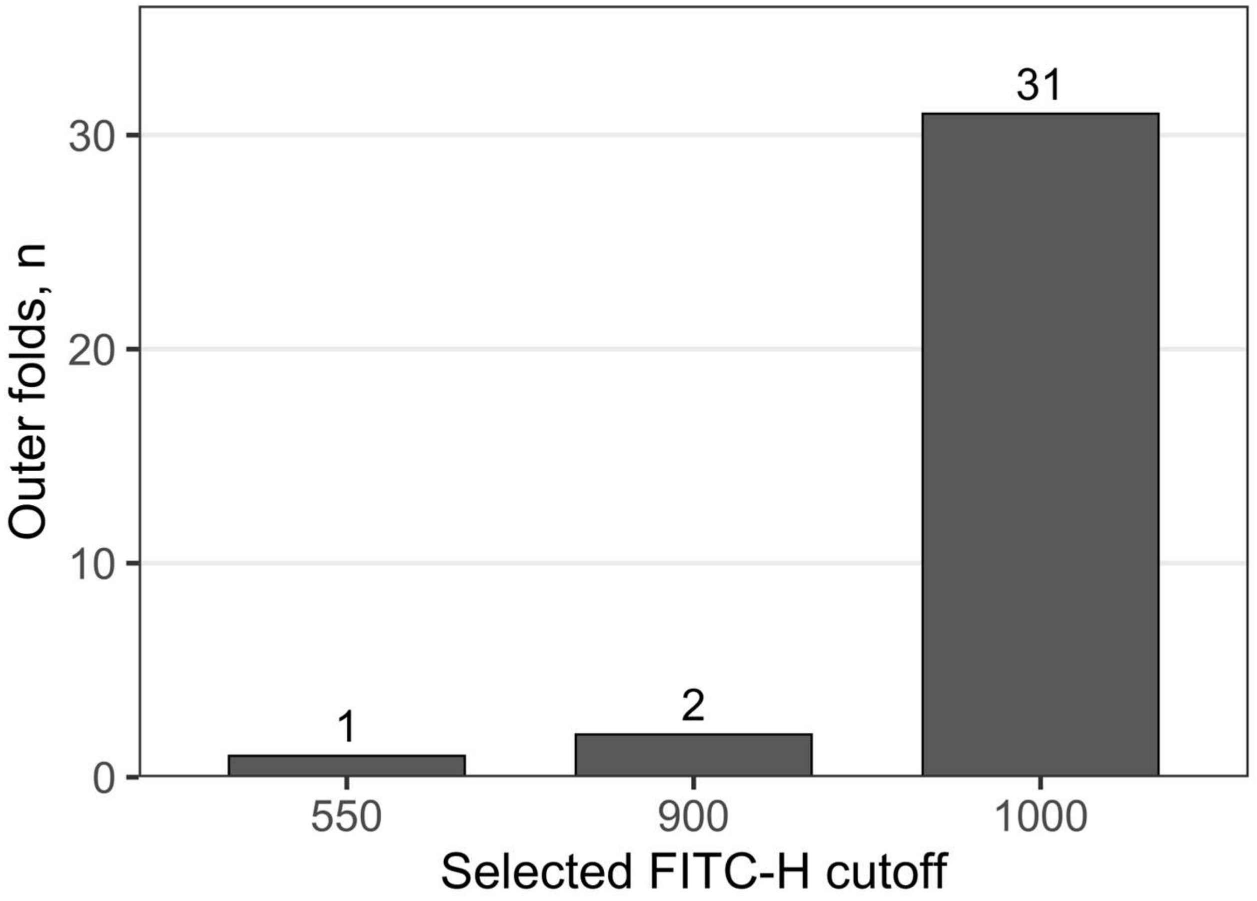


**Figure S5. Distribution of FITC-H cutoffs selected in post hoc nested LOOCV**
The selected cutoff was 1000 in 31 of 34 outer folds, 900 in two folds, and 550 in one fold.

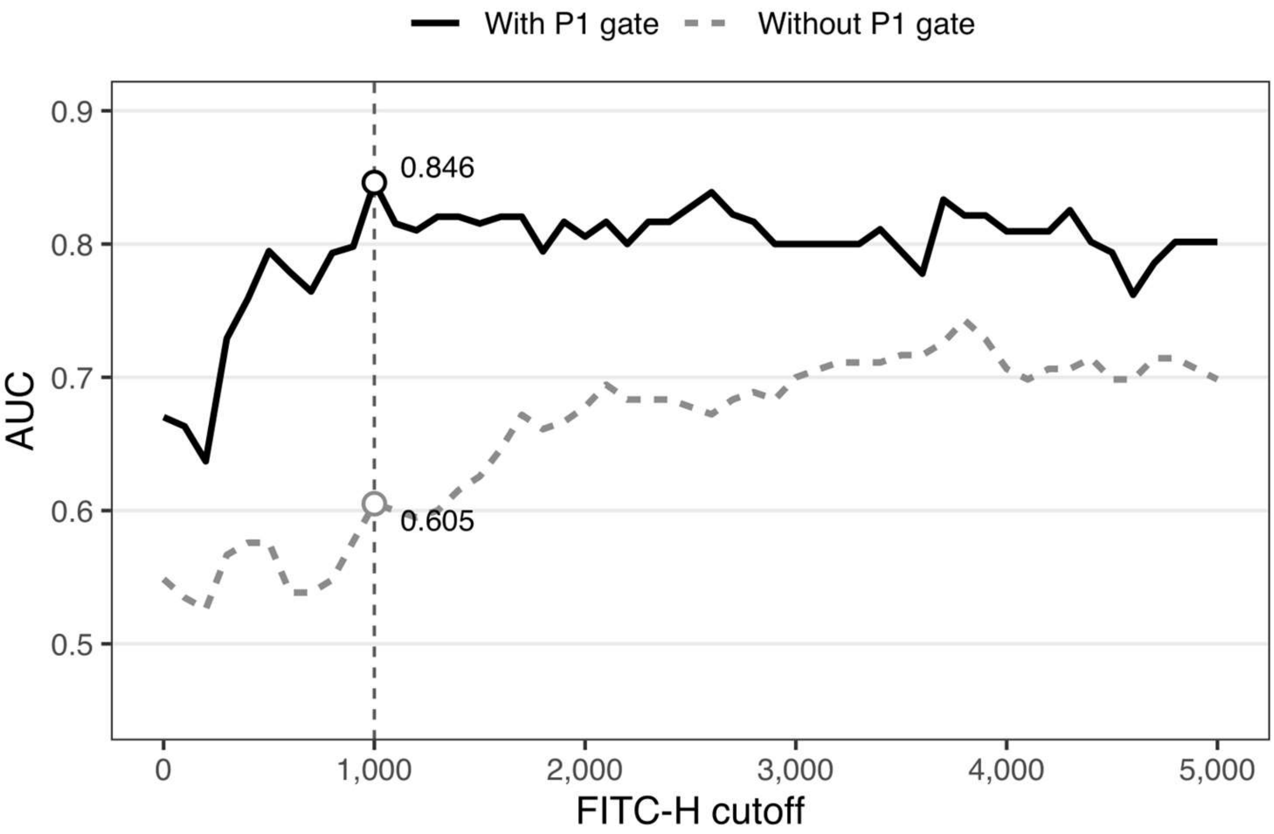


**Figure S6. Matched comparison of AUC across candidate FITC-H cutoffs with and without the P1 gate**

At each candidate cutoff, only samples with one or more retained events under both P1-gated and ungated analyses were included. Thus, the two conditions were compared using the same samples at each cutoff, although the matched sample set could change across cutoffs. Solid black and dashed gray lines indicate analyses with and without P1, respectively. At FITC-H = 1000, the AUC was 0.846 with P1 and 0.605 without P1.

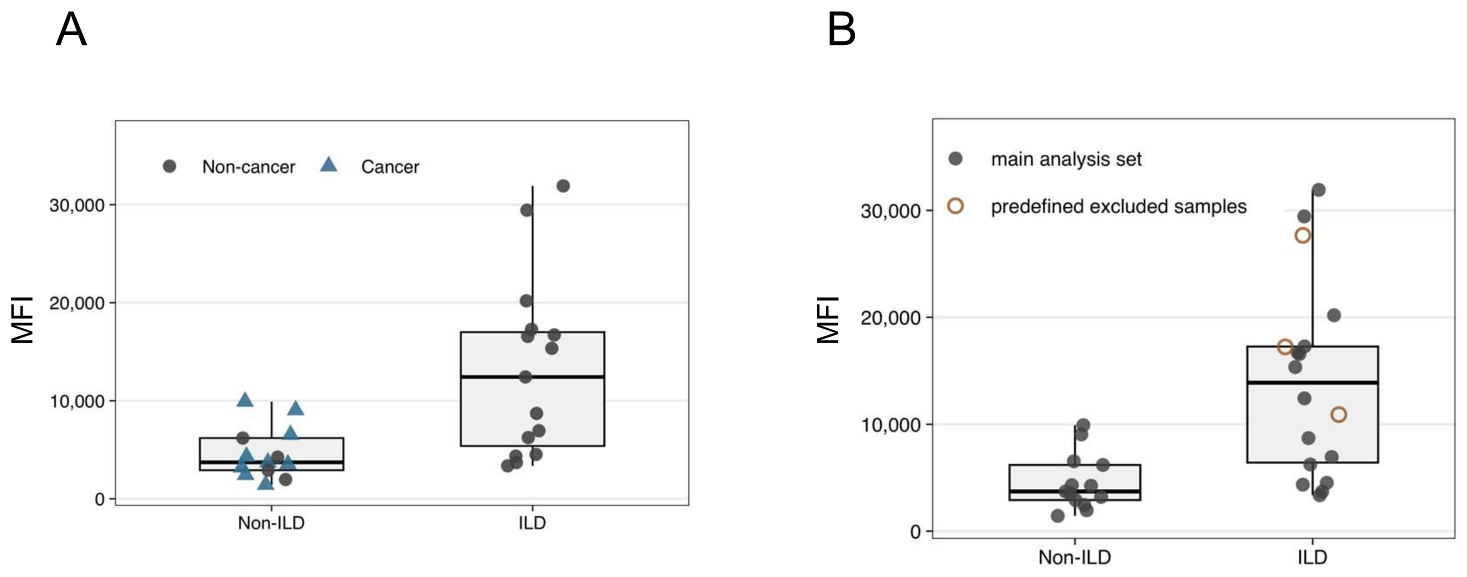


**Figure S7. MFI comparison between ILD and non-ILD groups in the primary and sensitivity analyses**

(A) MFI values in the ILD and non-ILD groups in the primary analysis set, stratified by cancer status. Non-cancer and cancer samples are indicated by circles and triangles, respectively. Blue triangles indicate samples from patients with thoracic malignancy. (B) Sensitivity analysis including the three samples that were excluded from the primary analysis. Filled symbols indicate samples included in the primary analysis, whereas open symbols indicate the predefined excluded samples. The sensitivity analysis included 31 (28 + 3 excluded) samples with events at or above FITC-H = 1000 (18 ILD and 13 non-ILD samples). The AUC was 0.872 (95% CI, 0.751–0.993), and the fixed-cutoff leave-one-out cross-validation (LOOCV) AUC was 0.803.

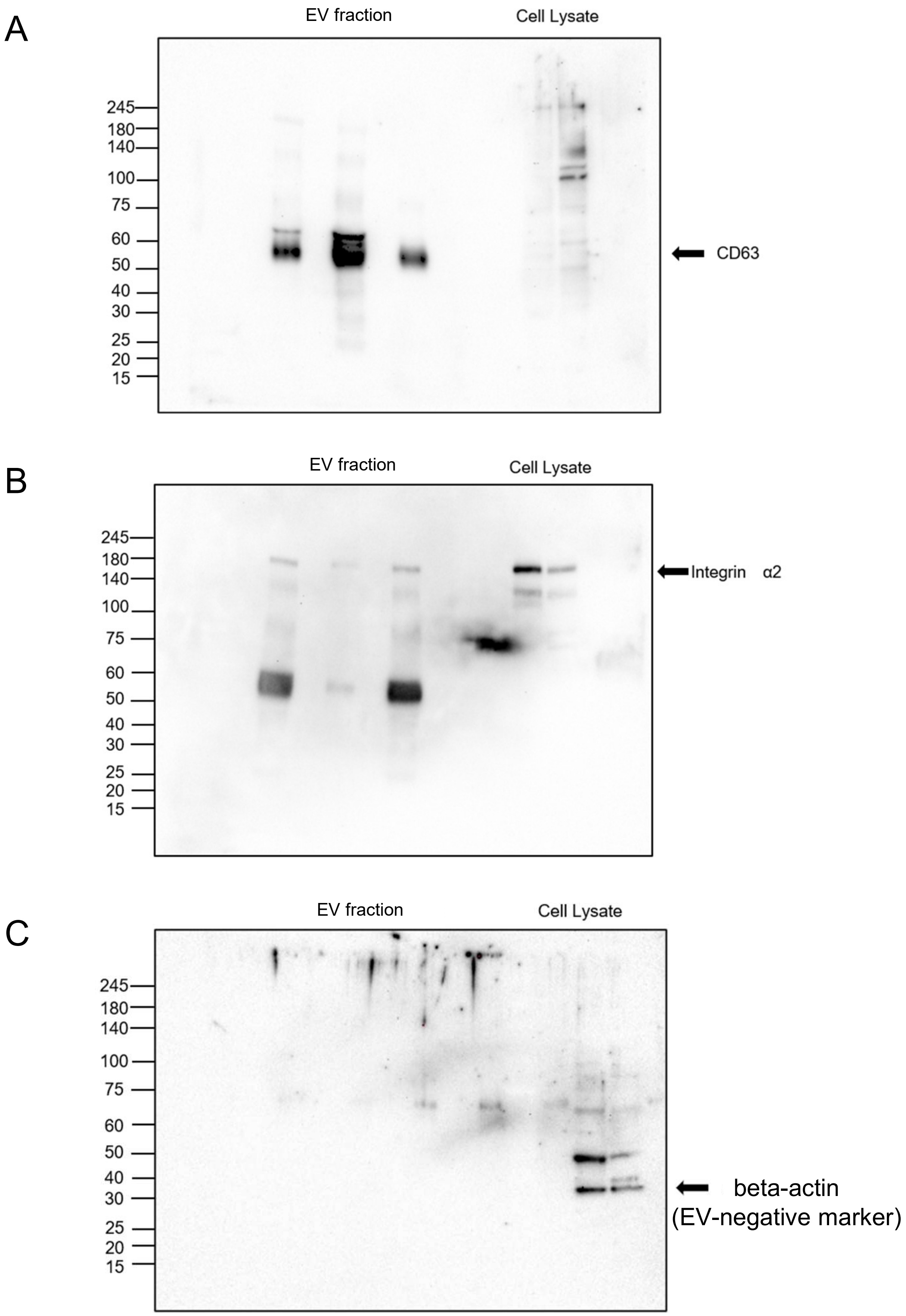


**Figure S8. Uncropped western blot images**

Uncropped western blot images corresponding to Fig. 2C. CD63, integrin alpha 2, and beta-actin are shown. Molecular mass markers are indicated in kDa.
